# Reduced Blood Flow in Deep White Matter During Hypercapnia Revealed by Multi-Delay pCASL in Healthy Young Adults

**DOI:** 10.64898/2026.06.01.729378

**Authors:** Yutong Lydia Sun, Nayana Menon, Xiaole Zachary Zhong, J. Jean Chen

## Abstract

The white matter (WM) cerebrovascular response remains poorly understood compared with grey matter (GM), partly due to technical challenges in perfusion quantification. Previous studies of cerebrovascular reactivity (CVR) have mostly been performed using BOLD MRI, and it is unclear to what extent they reflect changes in cerebral blood flow (CBF). In this work, we used multi-delay pseudo-continuous arterial spin labeling (pCASL) to quantify hypercapnia-induced CBF changes (ΔCBF) while accounting for regional variability in arterial transit time. Twenty-five healthy young adults underwent MRI during normocapnia and hypercapnia (inhalation of a 4% CO₂ gas mixture). Hypercapnia induced robust positive ΔCBF in cortical GM (26.7 ± 13.5%), superficial WM (17.2 ± 12.6%), periventricular regions (13.6 ± 10.6%), and subcortical GM (25.7 ± 14.1%) (all p < 0.0001). In contrast, deep WM exhibited a near-zero group-mean CBF response (1.0 ± 8.9%, p = 0.57), with 10 of 25 participants demonstrating negative ΔCBF. Negative CBF responses were consistently localized to the corona radiata, centrum semiovale, and optic radiation. Quality-control analyses showed that deep-WM ΔCBF estimates are robust, supporting the reliability of these findings. Moreover, across tissue compartments, higher baseline CBF was associated with reduced hypercapnic responsiveness, and deep-WM responses were strongly coupled with cortical GM responses across individuals. These results demonstrate that hypercapnia-induced cerebrovascular responses are highly heterogeneous across tissue depths and provide evidence that negative CBF responses can occur in healthy deep WM in the presence of vasodilation elsewhere. The findings challenge the assumption of uniformly positive perfusion responses during hypercapnia and support a potential role for flow redistribution arising from regional differences in vascular resistance and reserve capacity.

## 1 Introduction

The hemodynamic response of the white matter (WM) remains poorly understood. One avenue to investigating it is through quantifying cerebrovascular reactivity (CVR). Prior CVR works have focused predominantly on the grey matter (GM) (Driver et al., 2017; Ito et al., 2003; Nöth et al., 2008), partially because the GM has higher vascular density and baseline perfusion (Leenders et al., 1990), making CVR measurements more feasible. Most CVR studies have used blood-oxygen-level-dependent (BOLD) functional MRI (fMRI), although cerebral blood flow (CBF) is a more fundamental metric of the vascular response (Battisti-Charbonney et al., 2011; Halani et al., 2015). Although BOLD is theoretically multifactorial as it reflects changes in CBF and blood volume (CBV) among other factors (Kim & Ogawa, 2012), it has been assumed that the hypercapnia-evoked BOLD-CVR can be directly translated into corresponding changes in CBF (Ito et al., 2003). In particular, the unknown relationship between WM BOLD and CBF poses an extra challenge when quantifying WM CVR. Moreover, the WM has lower CBF and CBV, making it intrinsically harder to detect hemodynamic changes in the WM (de Riedmatten et al., 2025; Gawryluk et al., 2014). Furthermore, the WM exhibits longer and more heterogeneous arterial-transit times, making hemodynamic parameter estimation using non-invasive neuroimaging techniques even more challenging (van Gelderen et al. 2008; Gallichan and Jezzard 2009; van Osch et al. 2009). As a result, WM CVR has often been interpreted using physiological frameworks largely derived from GM observations (Sleight et al., 2022). Nevertheless, it is an unverified assumption that WM hemodynamics conform to those observed in the GM, as the WM has a distinct vascular architecture and oxygen metabolism (Leenders et al., 1990; Nonaka et al., 2003).

Hypercapnia is widely employed to probe CVR as a measure of hemodynamics (Liu et al., 2019). Hypercapnia refers to an increased arterial concentration of carbon dioxide (CO_2_) as a potent vasodilator stimulus which modulates the extracellular pH, leading to relaxation of vascular smooth muscle and reduced vascular resistance (Keyeux et al., 1995; Yoon et al., 2012). BOLD-based CVR provides high spatial resolution and whole-brain coverage, while remaining sensitive to changes in blood oxygenation associated with vascular responses. Using hypercapnia, past works have demonstrated a smaller BOLD-CVR in the WM compared to in the GM under the same level of vessel dilatory stimulus (Bhogal, 2021; Bhogal et al., 2015; Levine et al., 2025; Poublanc et al., 2015; Rostrup et al., 2000; Thomas et al., 2013). However, relatively few studies have directly validated CBF response to hypercapnic vasodilatory stimuli (Taneja et al., 2020). Collectively, there is a need for a more integrated assessment of hemodynamics in each of WM and GM using quantitative, flow-sensitive MR techniques.

Early works using pulsed arterial-spin labeling (ASL) reported almost exclusively on GM (Driver et al., 2017; Nöth et al., 2008). Very few early works showed that WM exhibited a temporally delayed and attenuated CBF response compared to GM when measured (Rostrup et al., 2000). Pseudo-continuous arterial-spin labeling (pCASL) has become the de facto standard ASL technique (Pinto et al., 2023; Thomas et al., 2013; Zhou et al., 2015). However, most prior works using pCASL applied single post-labeling delays (PLD) of 1.2-1.5 s (Thomas et al., 2013; Zhou et al., 2015), which is shorter than the typical average WM ATT (Juttukonda et al., 2021). As a result, labeled blood water may not have fully arrived in distal WM tissue at the time of image acquisition, potentially leading to systematic CBF underestimation and reduced interpretability of WM hemodynamics. Incorporating multiple PLDs in pCASL can improve CBF quantification by accounting for inter-regional (i.e. WM vs. GM) ATT variability (Alsop et al., 2015). This is particularly important for this study considering the long and heterogeneous transit delays in the WM (Gallichan & Jezzard, 2009; van Osch et al., 2009). The heterogeneity in acquisition strategies in earlier works also complicates cross-study comparisons.

In this study, we assessed hemodynamics in WM and compared them with those in GM using a hypercapnia challenge combined with multi-delay pCASL. Our work aims to provide a more comprehensive assessment of regional CBF response to CO_2_ in the WM. To verify the reliability of WM CBF measurements, we additionally performed a dedicated quality control analysis evaluating the robustness of CBF estimates in deep WM.

## 2 Methods

### 2.1 Participants

Thirty young adults without a diagnosis of neurological, psychological or major physical diseases were recruited. All participants provided informed written consent in accordance with the Research Ethics Board approval of the Baycrest Academy for Research and Education. Of the 30 participants, 27 completed all MRI scan sessions involving breathing tasks. 2 participants were excluded at the analysis stage due to motion artifacts (bulk head motion), resulting in a final sample of 25 participants (13 female, 12 male, mean age of 25 ± 4 years).

### 2.2 Gas Inhalation Experiment

Hypercapnia was induced by having participants inhale a gas mixture containing 4% CO_2_, 19% O_2_, and 77% N_2_ via a custom-built gas delivery system designed to ensure stable and reproducible gas administration. The CO_2_ level was modest in order to minimize the likelihood of confounding neuronal activation (Chen & Pike, 2010), and each hypercapnic session lasted approximately 6-7 minutes (6:08 min scan + ∼30 second pre-scan CO_2_). The end-tidal partial pressure of CO_2_ (PETCO_2_), breathing rate, and heart rate were monitored in real time to confirm the reach of hypercapnic states as well as full recovery to post-hypercapnic baseline after the scan ends. For a more detailed explanation see Supplementary Figure 1.

### 2.3 MRI Acquisitions

MRI scans were performed on a Siemens Prisma 3 Tesla MRI scanner using a 32-channel receive-only head coil. High-resolution T1-weighted structural images were acquired using magnetization-prepared rapid gradient-echo (MPRAGE) with TR = 2 s, TE = 2.85 ms, TI = 950 ms, 0.8 mm isotropic resolution. pCASL data were acquired using a 2D multi-band echo-planar imaging (MB-EPI) readout under both hypercapnic and normocapnic conditions, with the following parameters: TR = 3.95 s, TE = 19 ms, in-plane resolution = 2.5×2.5 mm^2^, slice thickness = 2.81 mm. Considering that CBF temporal dynamics differ between hypercapnic and normocapnic conditions, pCASL data were collected on multiple PLDs to prevent CBF underestimation (Pinto et al., 2023). Tag–control pairs were acquired sequentially across PLDs, with 6 pairs collected at PLD = 0.4 s, followed by 6 pairs at 0.9 s, 6 pairs at 1.4 s, 10 pairs at 1.9 s, and 14 pairs at 2.4 s. The labeling duration was 1.5 s. A pair of field maps were also acquired on the reversed phase-encoding directions. The pCASL acquisition time was 6:08 minutes.

### 2.4 Data Processing

T1-weighted structural images were processed with FreeSurfer’s *recon-all* pipeline (v7.4.1), which includes skull stripping, intensity normalization, tissue segmentation and cortical and subcortical parcellation (Dale et al., 1999; Desikan et al., 2006; Fischl et al., 2002). pCASL data were first corrected for susceptibility-induced distortions using FSL’s *topup* (Andersson et al., 2003). Quantitative CBF maps in units of mL/100g/min were then generated using the general kinetic model implemented in *oxford_asl* (BASIL, v4.0.29) *within FSL (v6.0.7.10)* (Chappell et al., 2020), incorporating tissue-specific ATT estimation and partial volume effect (PVE) correction.

### 2.5 Statistical Analysis — CO_2_-induced CBF Change

Percent CBF change (%ΔCBF) due to hypercapnia was calculated voxelwise and registered to the corresponding T1-weight space using an affine transformation implemented in FSL FLIRT with nearest-neighbour interpolation (Jenkinson et al., 2002). Group-level voxelwise analysis was performed using a one-sample t-test, with statistical significance defined using threshold-free cluster enhancement (TFCE) with 5000 permutations and family-wise error (FWE)-corrected p < 0.05. Structural data were further used to segment the brain into five regions-of-interest (ROIs) based on their tissue depth: cortical GM, superficial WM, deep WM, periventricular region, and subcortical GM (Figure 1). This segmentation was performed in each participant’s native space using the automatic cortical parcellation (aparc) and automatic segmentation volume (aseg) outputs generated by recon-all, from which cortical GM, whole WM, ventricular, and subcortical GM masks were obtained. Tissue-depth-specific ROIs were subsequently generated using morphological operations: the deep WM mask was created by eroding the whole-WM mask by four voxels, the superficial WM mask was defined as the remaining WM after subtraction of the deep WM mask, and the periventricular mask was generated by dilating the ventricular mask by two voxels. Regional average ΔCBF, in both percent and absolute units (mL/100g/min), was calculated and evaluated against the regional average baseline CBF (CBF_0_) in corresponding ROIs. Note that all statistical analyses were carried out on voxels with %ΔCBF values falling within the range of ±99.9% to ensure robust and physiologically plausible values (Kety & Schmidt, 1948).

**Figure 1.**
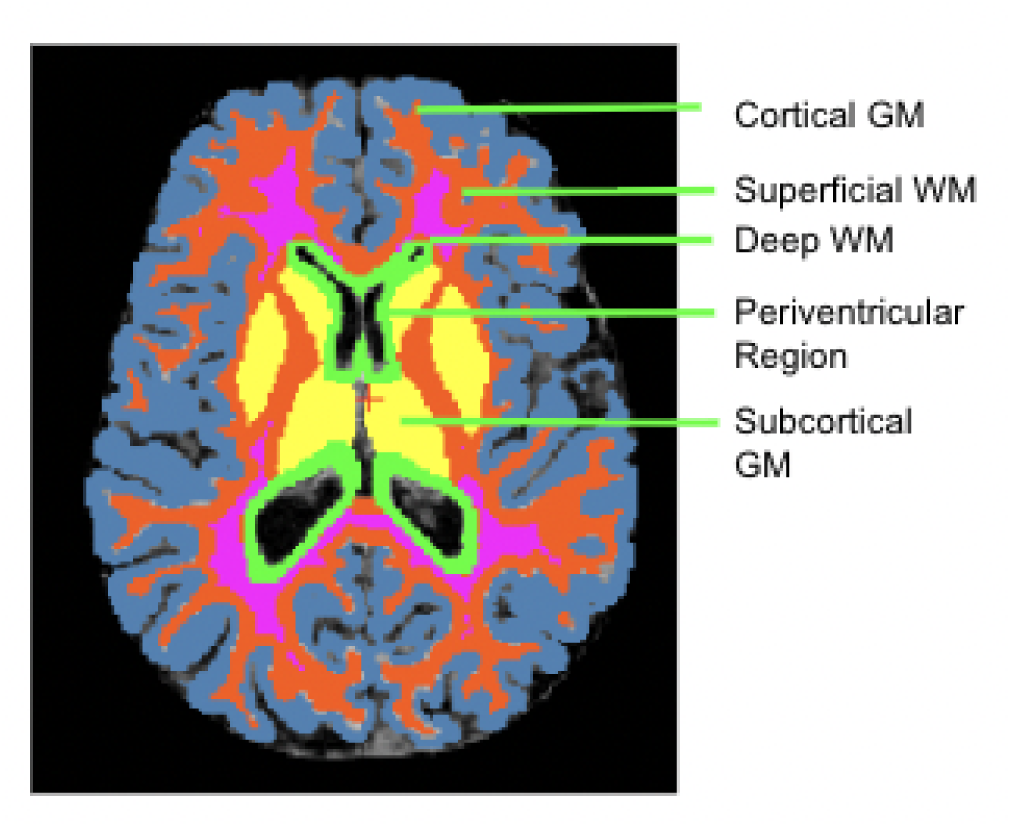
Brain map illustrating the definition of tissue-depth-based regions-of-interest (ROIs) used for regional CBF and ΔCBF analysis. A T1-weighted structural image in one representative participant’s native space processed with FreeSurfer were segmented into cortical grey matter (GM) and white matter (WM), and further partitioned into depth-dependent ROIs based on distance from the cortical surface, including cortical GM, superficial WM, deep WM, periventricular region, and subcortical GM.

### 2.6 Quality Control (QC) of WM CBF Response Quantification

To assess the reliability of CBF measures in WM, we investigated the effect of the number of measurements on WM %ΔCBF estimates. To that end, a total of 42 pCASL tag-control pairs were first reorganized into six PLD sets, with each set containing one tag-control pair at each of the five PLDs. Hypercapnic and normocapnic CBF, as well as CO_2_-induced %ΔCBF and its corresponding t-statistics from a one-sample t-test of %ΔCBF (to assess how it significantly differs from zero), were computed using cumulatively increasing numbers of tag-control pairs in PLD sets. For a more detailed explanation see Supplementary Figure 2.

Since a greater number of pairs were acquired at longer PLDs (1.9 and 2.4 s), the influence of additional long-PLD data was further examined by incorporating these additional bonus pairs with the 6 complete PLD sets. We hypothesized that the 6 complete PLD sets plus the bonus long-PLD pairs permit stable CBF and %ΔCBF estimates in the WM.

## 3 Results

Successful induction of the hypercapnic state was confirmed by changes in steady-state PETCO_2_. Mean (±SD) PETCO_2_ across participants increased from 4.0% (± 0.6%) at baseline to 5.0% (± 0.4%) during hypercapnia (p < 0.0001).

As shown in Figure 2, hypercapnia induced widespread CBF increases in cortical and subcortical GM. In contrast, CBF responses to hypercapnia in WM regions were heterogeneous, with both positive and negative responses consistently present across participants but varying in magnitude. Figure 2a shows the CBF_0_ and %ΔCBF maps from a participant with a representative negative CBF response in deep WM, while Figure 2b shows those of another participant with a representative positive %ΔCBF in deep WM. Notably, negative hypercapnic CBF responses were still present in localized deep-WM regions even when the regional mean %ΔCBF was positive (Figure 2b).

**Figure 2.**
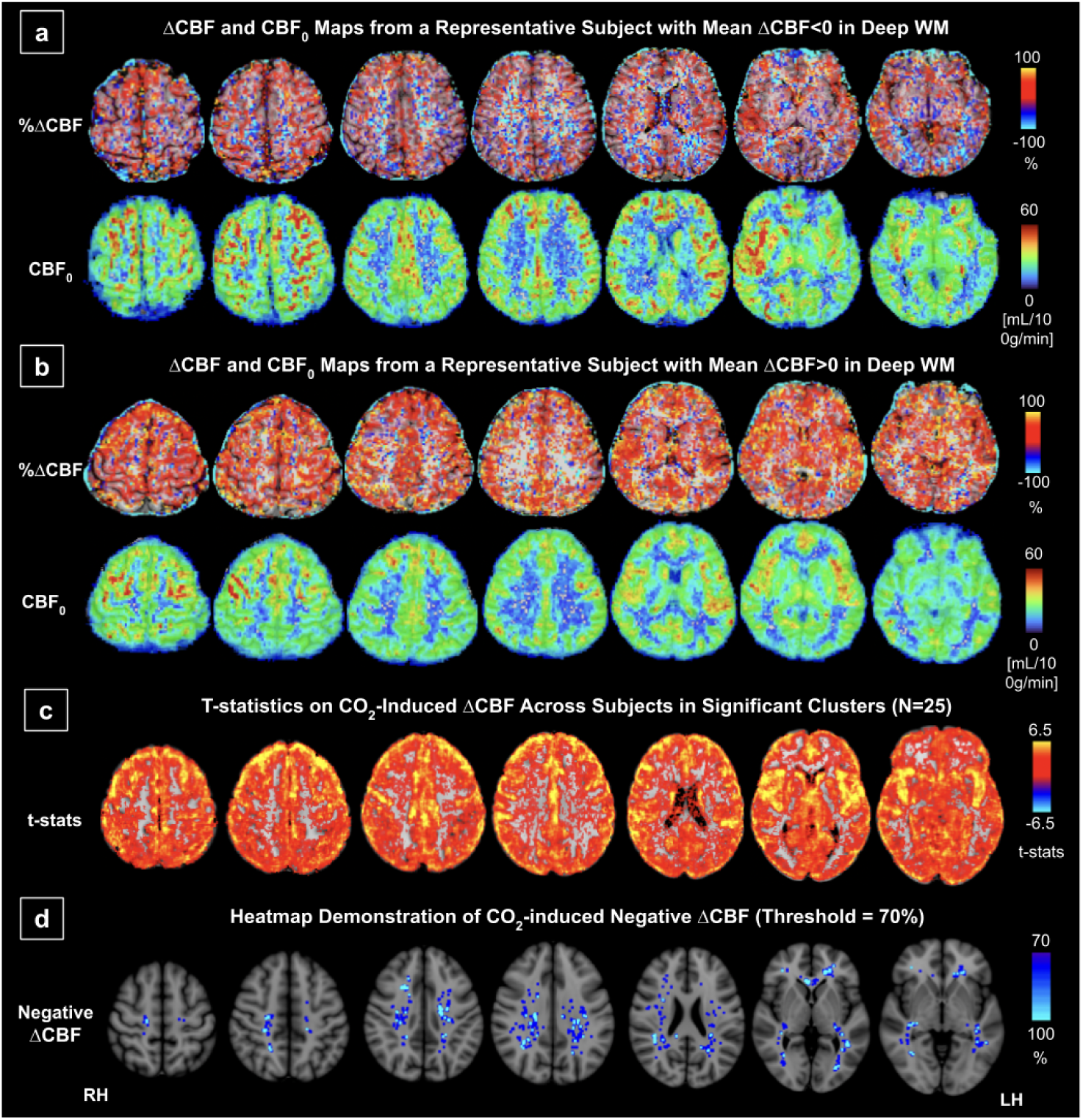
Baseline CBF (CBF_0_) and CO_2_-induced percent CBF change (%ΔCBF) maps, along with group-level voxelwise statistics. (a) Axial slices in the participant’s native space for %ΔCBF (top row) CBF_0_ (bottom row) from a representative participant exhibiting a net negative mean %ΔCBF within deep WM. (b) Axial slices in the participant’s native space for %ΔCBF (top row) CBF_0_ (bottom row) from a representative participant exhibiting a net positive mean %ΔCBF within deep WM. Blue voxels are present in both representative participants but differ in spatial extent and magnitude. (c) group-level voxelwise t-statistics from a one-sample t-test of CO_2_-induced %ΔCBF against zero, overlaid on structural images in MNI152 space, thresholded using threshold-free cluster enhancement; red-to-yellow indicates statistically significant CO_2_ effect and greyscale indicates absence of statistically significant effects at group level. (d) Heatmap generated on a subset of participants with negative average deep-WM %ΔCBF (n=10) (see Figure 2), highlighting voxels where ≥70% of participants exhibited negative CO_2_-induced %ΔCBF, dilated with maximum filtering, overlaid on structural images in MNI152 space.

Voxelwise one-sample t-tests identified significant positive CO_2_-induced ΔCBF predominantly in cortical GM (Figure 2c). A gradual attenuation of the positive CBF response was observed with increasing tissue depth. Spatially, positive CBF responses were more pronounced in the bilateral frontal and lateral regions of the brain. Negative CBF responses that were observed in deep WM at the individual level did not reach voxelwise significance at the group level. Nevertheless, among those that exhibited negative ΔCBF in the deep WM, spatially consistent regions can be seen in Figure 2d (regions that show a negative ΔCBF in over 70% of the participants with negative average deep-WM ΔCBF). These responses were primarily localized in the corona radiata, centrum semiovale, and optic radiation.

Group-level ROI analysis (Figure 3) showed significant CBF increase of 26.7%±13.5% (p < 0.0001), 17.2%±12.6% (p < 0.0001), 13.6%±10.6% (p < 0.0001), 25.7%±14.1% (p < 0.0001) in cortical GM, superficial WM, periventricular region, and subcortical GM, respectively. In contrast, deep WM showed a mean %ΔCBF of 1.0%±8.9% (p=0.57). This low value is representative of the low reactivity of the WM, but can also be attributed to inter-participant differences, with 10 of 25 participants exhibiting a net average %ΔCBF below zero (Figure 3). Significant inter-ROI differences (p < 0.05) were detected except for between cortical GM and subcortical GM, as well as between superficial WM and periventricular regions.

**Figure 3.**
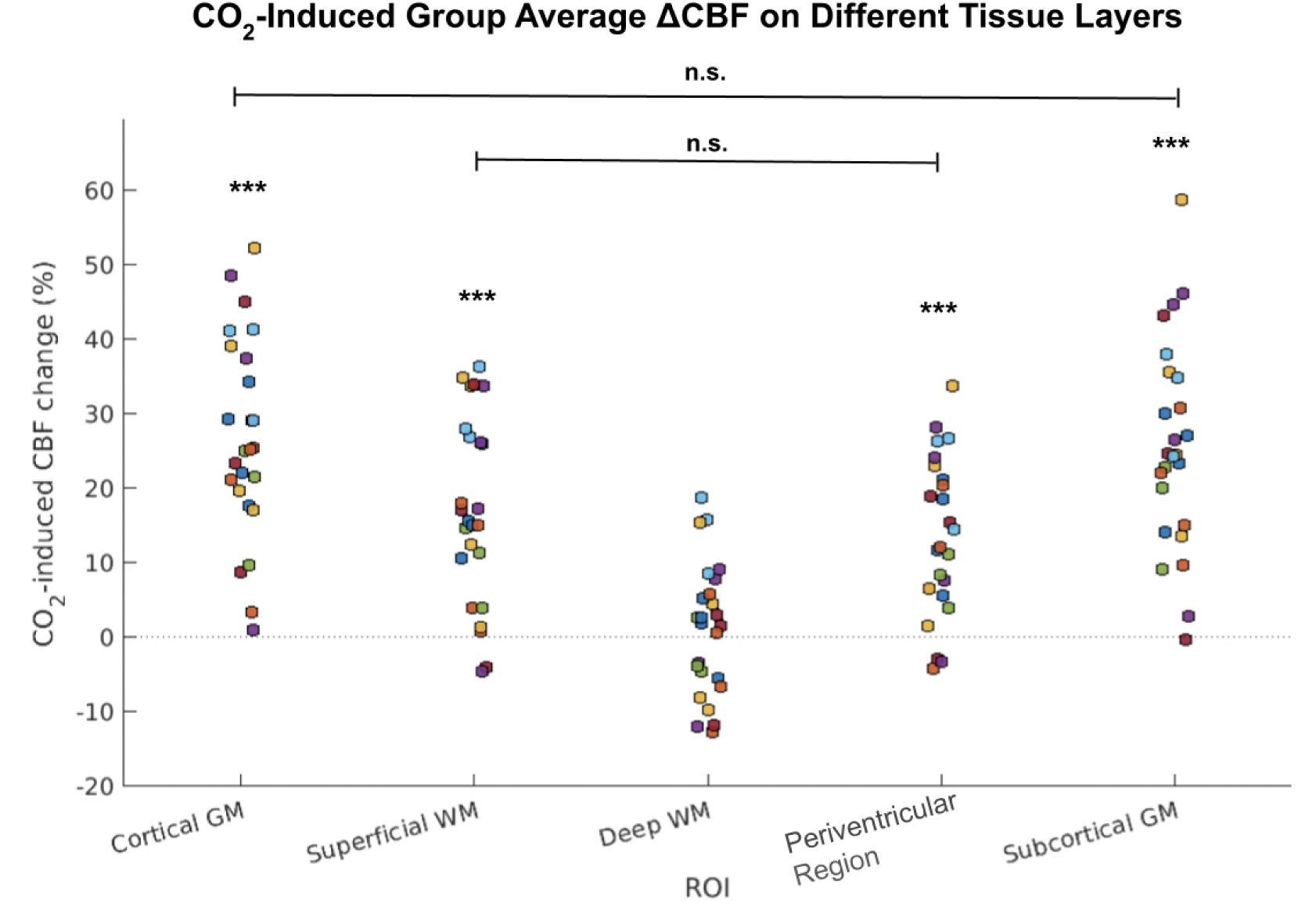
Mean CO_2_-induced percent CBF change (%ΔCBF) across the tissue-depth-defined ROIs. Each data point represents the mean %ΔCBF from an individual participant within a given ROI, including cortical GM, superficial WM, deep WM, periventricular regions, and subcortical GM. Inter-ROI comparisons were done by paired t-test with Bonferroni correction: ROIs labeled with n.s. are not significantly different from each other, while all other comparisons are significant. Significant CBF increases were observed in all ROIs except deep WM, which showed a near-zero mean %ΔCBF with substantial inter-participant variability on both positive (n = 15) and negative (n = 10) responses.

To ensure the reliability of these WM ΔCBF results, especially the negative ΔCBF findings, QC analyses were carried out on participants with negative average ΔCBF in deep WM to better assess whether the observed negative CBF response could be due to acquisition limitations (Gallichan & Jezzard, 2009; van Osch et al., 2009). Figure 4 shows the group averages of CBF under the two gas conditions, as well as the CO_2_-induced %ΔCBF, and the corresponding t-statistics. We observed that CBF estimates under both gas conditions, as well as the CO_2_-induced %ΔCBF and its corresponding t-statistics, varied as additional PLD sets were included in the kinetic-model fitting. In particular, as more long-PLD tag–control pairs were incorporated, CBF, %ΔCBF, and t-statistics plot increasingly stabilized, suggesting that the estimates become relatively stable in deep white matter. With all 6 complete PLD sets and 12 additional long-PLD pairs, we can observe robust non-zero estimates of %ΔCBF and t-statistics.

**Figure 4.**
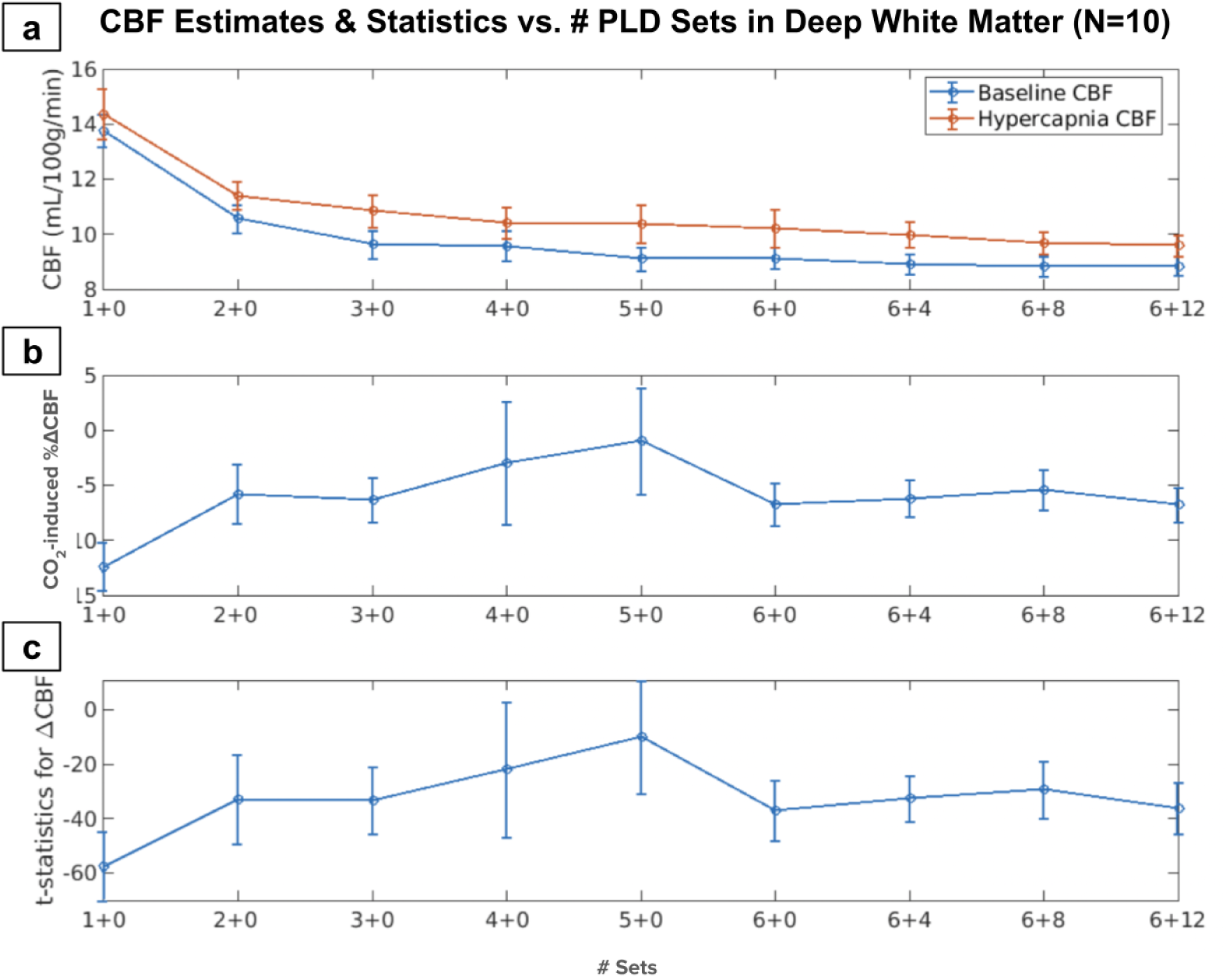
Deep WM CBF estimations and statistics as a function of the number of pCASL tag-control pairs, evaluated in participants with negative average deep-WM %ΔCBF (n=10; see. Figure 2). The x-axis denotes the different amounts of data used for the CBF estimation. Specifically, labels “1+0” through “6+0” represent the number of complete 5-PLD sets, where one complete PLD set consists of one tag–control pair acquired at each PLD. Labels beyond this point include additional long-PLD pairs: “6+4” indicates 6 complete PLD sets plus 4 additional long-PLD tag-control pairs (2 at PLD = 1.9 s and 2 at PLD = 2.4 s); “6+8” indicates 6 complete PLD sets plus 8 additional tag-control pairs at longer PLDs (4 at 1.9 s and 4 at 2.4 s); and “6+12” indicates 6 complete PLD sets plus 12 additional long-PLD tag-control pairs (4 at 1.9 s and 8 at 2.4 s). The error bars represent the group-wise standard deviation of the regional means across participants. (a) Mean baseline CBF (normocapnia) and hypercapnic CBF (mL/100g/min) across participants estimated in deep WM as progressively more pair sets and long-PLD pairs were incorporated into the kinetic model fitting. (b) Corresponding CO_2_-induced percent CBF change (%ΔCBF) in deep WM. (c) t-statistics from a one-sample t-test of %ΔCBF against zero.

To understand the determinants underlying the occurrence of both positive and negative ΔCBF, the relationship between CO_2_-induced %ΔCBF and CBF_0_ was investigated on tissue-depth-defined ROIs. Percent ΔCBF exhibits a significant negative association with CBF_0_ across most tissue ROIs, except for deep WM (Figure 5a). The same analysis was conducted using quantitative ΔCBF in mL/100g/min to rule out possible bias from mathematical coupling between %ΔCBF and CBF_0_. Quantitative ΔCBF shows weaker and more region-specific negative relationships with CBF_0_, with significant associations primarily observed in deep WM (Figure 5b). Moreover, strong positive associations were observed between deep WM ΔCBF and cortical GM ΔCBF for both percent and absolute measures (Figure 5c–d), indicating coordinated perfusion responses across tissue depths within participants.

**Figure 5.**
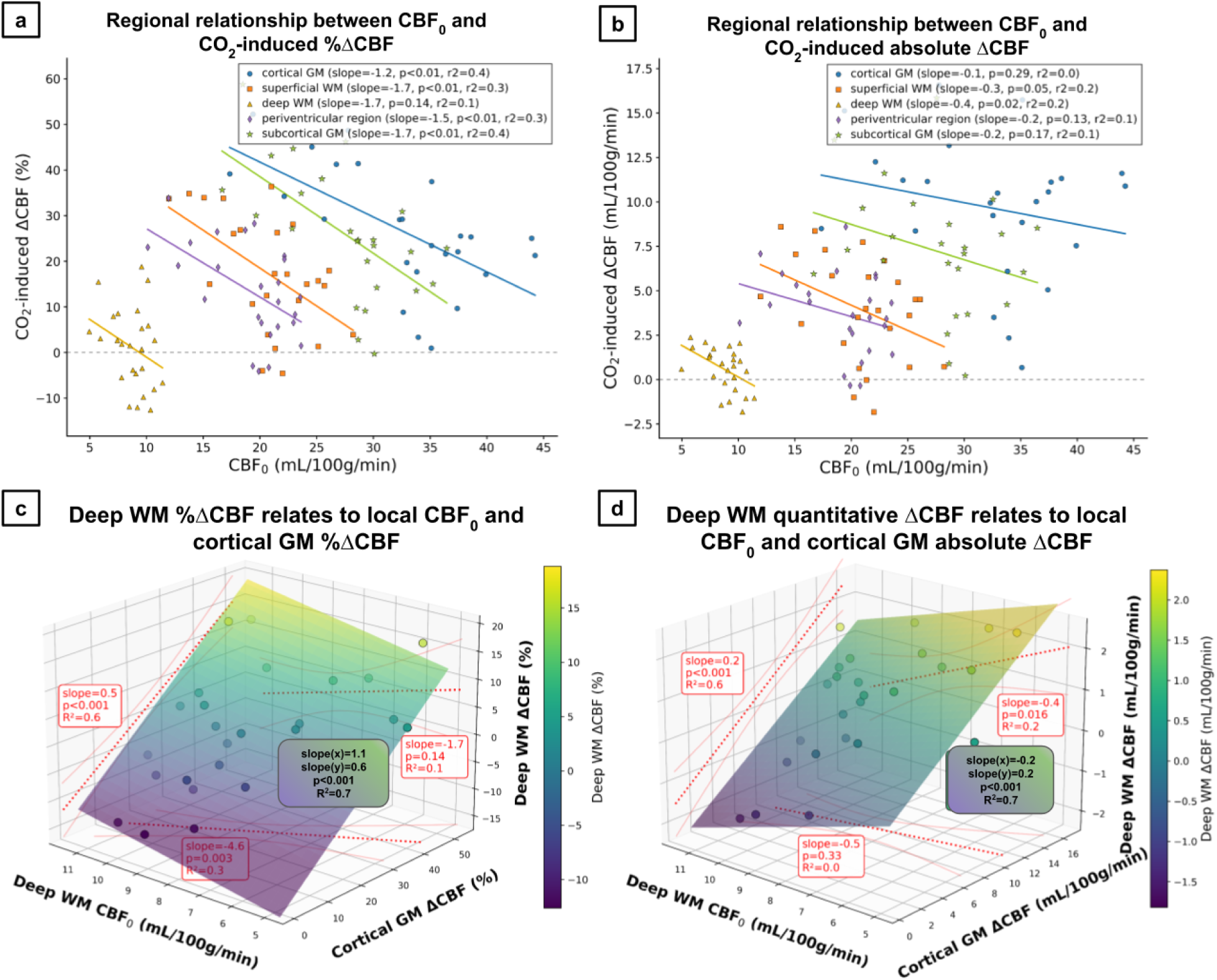
Regional relationships between baseline CBF (CBF_0_) and CO_2_-induced percent ΔCBF (left) and quantitative ΔCBF (right). (a) Relationship between CBF₀ and CO₂-induced percent ΔCBF across five tissue-depth-defined ROIs: cortical grey matter (cGM), superficial white matter (sWM), deep WM (dWM), periventricular region (pvR), and subcortical GM (scGM), distinguished by colour and marker type. Each data point represents the mean values from an individual participant within a given ROI. Solid lines indicate least-squares linear fits for each region. Corresponding slope, p-value, and coefficient of determination (R²) for each regression are reported in the legends. (b) Same analysis as in (a) using quantitative ΔCBF in mL/100g/min. (c) Three-dimensional relationship illustrating how dWM percent ΔCBF relates jointly to local CBF_0_ and cGM percent ΔCBF. Each point represents a participant, and the colored plane shows the fitted multiple linear regression surface. Data points colour represents dWM percent ΔCBF values. Regression coefficients (slopes), p-values, and R² values for the multivariable model, as well as for the pairwise relationships, are annotated within each panel. (d) Same analysis as in (c) using quantitative ΔCBF in mL/100g/min. Together, these results demonstrate the inter-regional coherence of CO_2_-induced perfusion responses and region-dependent coupling between perfusion responses and baseline perfusion.

## 4 Discussion

The WM vascular response to hypercapnic vasodilatory stimuli has been commonly assumed to follow that of the GM (Taneja et al., 2020). However, in this work, by examining CBF changes under a hypercapnic challenge with ATT explicitly accounted for, we observed that deep WM can exhibit negative CBF responses during hypercapnia (Figure 2a, 2b). This response pattern contrasts with the robust positive CBF response consistently observed in GM and challenges the conventional assumption that hypercapnia induces CBF increases throughout the brain. We will elaborate on these findings as follows.

### 4.1 CBF Quantification in the White Matter

There are long-standing challenges in measuring WM CBF with ASL (Gallichan & Jezzard, 2009; van Gelderen et al., 2008; van Osch et al., 2009). Compared with GM, WM regions are characterized by longer ATT as they are more distal from major arterial sources (van Gelderen et al., 2008). Although the PLDs employed in this study (0.4 to 2.4 s) are generally considered sufficient for capturing labeled blood arrival in most brain regions in young healthy individuals (Alsop et al., 2015), ATT in very deep WM regions may extend beyond 3 s (van Gelderen et al., 2008). The prolonged ATT and a subsequent greater bolus dispersion likely result in fluctuating CBF estimations (Gallichan & Jezzard, 2009). While prior studies have appropriately emphasized factors such as prolonged and heterogeneous arterial transit times, bolus dispersion, and reduced labeling efficiency may compromise WM perfusion quantification (Gallichan & Jezzard, 2009; van Gelderen et al., 2008; van Osch et al., 2009), these findings were largely derived from single-PLD acquisitions, and in some cases using earlier ASL implementations with lower SNR. In contrast, our study employed a multi-delay pCASL protocol with extended PLDs and additional long-PLD sampling, which allowed improved characterization of delayed inflow and reduced sensitivity to ATT-related biases. We performed detailed QC to test that, with a particular focus on the robustness of the negative ΔCBF findings. In this QC work, as seen in Figure 4a, the observed CBF estimates reached a stable state with increasing numbers of tag-control pairs, which supports the interpretation that deep WM CBF estimates obtained using our study are robust (Figure 4a). More importantly, because the primary focus of this work is the CO₂-induced ΔCBF, we also evaluated the stability of the ΔCBF estimates in the WM. As seen in Figures 4b and 4c, as the number of tag-control pairs increased, particularly with the inclusion of long-PLD pairs, both %ΔCBF and associated *t*-statistics demonstrated variations before stabilizing at non-zero values, suggesting reliable detectability of CO_2_-induced ΔCBF.

### 4.2 CBF Response to Hypercapnia in the Grey Matter

In line with the positive CVR identified in many previous BOLD-based CVR studies (Poublanc et al., 2015; Rostrup et al., 2000; Yezhuvath et al., 2009), our research found a robust and widespread CO_2_-induced CBF increase in cortical and subcortical GM regions (Figure 3). Unlike the more commonly used BOLD-CVR approach, which reflects a complex combination of cerebral blood flow, cerebral blood volume, oxygen metabolism, and baseline vascular state, ASL-derived CBF provides a more direct measure of the underlying vascular perfusion response to CO_2_ (Ainslie and Duffin 2009; Driver et al. 2017).

GM exhibits a stronger CBF response to CO_2_ than WM, consistent with earlier investigation using pASL and BOLD-CVR (Levine et al., 2025; Rostrup et al., 2000). The stronger vascular response in GM is likely attributable to its substantially higher vascular density compared with WM, leading to the potential for a greater reduction of vascular resistance during hypercapnia (Ainslie and Duffin 2009). In particular, the higher number of low-resistance vessels, including arterioles and precapillary vessels, may facilitate even stronger CO_2_-induced vasodilation and greater perfusion modulation in GM (Dabertrand et al., 2012; Levine et al., 2025). Despite the anatomical differences in arterial supply between cortical GM and subcortical GM, the magnitude and direction of hypercapnic responses were comparable. Cortical GM receives blood supply from pial and penetrating arteries (Agarwal & Carare, 2020), whereas subcortical GM is supplied by deep perforating arteries branching directly from major vessels (Vogels et al., 2021). These architectural differences may contribute to a shorter response time in subcortical GM than cortical GM, yet both regions are highly vascularized and dominated by arterioles that demonstrate similar perfusion modulation, suggesting broadly comparable vasodilatory behaviour.

### 4.3 CBF Response to Hypercapnia in the White Matter

Overall, WM demonstrated weaker hypercapnic perfusion responses than GM, with superficial WM generally exhibiting positive ΔCBF, whereas deep WM showed markedly reduced or even negative CO_2_-induced CBF responses in young healthy participants (Figure 2a, 2b, 2d, 3). These findings are consistent with the previous studies reporting reduced positive WM perfusion responses during hypercapnia compared with GM (Bhogal et al., 2015; Hartmann et al., 2021; Rostrup et al., 2000), as well as studies reporting negative hypercapnic responses in deep WM using both ASL-derived CBF (Mandell et al., 2008) and BOLD-CVR approaches (Arteaga et al., 2014; Bhogal et al., 2015; Thomas et al., 2013). It is important to note that because BOLD-CVR does not represent a direct measure of vascular perfusion response, negative BOLD-CVR responses may occur even in the absence of true CBF reduction, for example due to increased deoxygenated CBV during hypercapnia. In the present study, by using multi-delay ASL, we provide additional evidence that true negative responses can occur in healthy deep WM. To our knowledge, few prior studies have reported CBF responses that are predominantly localized to WM. Mandell et al. (2008) previously demonstrated selective negative hypercapnic CBF responses in the WM using single-delay FAIR pulsed ASL at 1.5T. However, the majority of negative CBF responses in their findings were found in what appears to be periventricular WM, with a different interpretation than the deep WM response. The use of pulsed ASL with a single PLD is also prone to insufficient capturing of WM perfusion and sensitivity to CO_2_-led ATT change (Alsop et al., 2015; Woods et al., 2024). In the present work, by using multi-delay pCASL with ATT correction at 3T, which is robust against both of these effects, we demonstrate clear negative CBF responses that are in deep WM. Furthermore, we uncover the inter-participant variability of these effects, formally connecting them to the vascular-resistance and reserve-capacity theory proposed by Duffin et al. (2018), where the negative CBF response is likely due to blood flow redistribution governed by inter-regional differences in vascular responsiveness and vascular reserve. These mechanisms will be further discussed in the following sections (see *Potential Mechanisms for Negative CBF Response in WM*).

While negative BOLD-CVR responses have been more frequently reported in deep WM (Arteaga et al., 2014; Bhogal et al., 2015; Thomas et al., 2013), the primary physiological mechanism underlying these responses remains debated. In principle, the CO_2_-induced decrease in BOLD-CVR in WM could be theoretically attributed to decreased CBF, increased CBV, increased oxygen metabolism and decreased partial pressure of oxygen (Kim & Ogawa, 2012). Ito et al. suggested that BOLD-CVR is predominantly flow-driven, whereas Arteaga et al. observed limited correspondence between negative BOLD-CVR and negative CBF responses (Alsop et al., 2015; Ito et al., 2003). The present findings demonstrate that negative CBF responses may occur even in healthy WM, implying that the net BOLD response likely reflects the balance between positive and negative vascular effects rather than a single dominant mechanism.

The absence of a significant negative CBF response at the group level (Figure 2c) is likely attributable to the substantial inter-participant variability in the extent and localization of CBF changes. This, however, does not diminish the physiological relevance of the observed negative CBF responses, as the thresholded heatmap demonstrates consistent localization of negative CBF responses across participants in deep WM, with nearly half of the participants exhibiting this behaviour (Figure 2d). Moreover, the magnitude of negative CBF responses may be underestimated in the present study due to venous contributions to the pCASL signal. Since the standard kinetic model does not explicitly account for venous outflow, the venous signal could be misinterpreted as blood exchanges between compartments and therefore bias CBF upward. This effect could be amplified in hypercapnic conditions due to reduced ATT and faster capillary-to-venous passage. In addition, hypercapnia was found to be related to enlarged venous radius, which may bias the measure even further and consequently shifting ΔCBF toward more positive values (Shen et al., 2013). Although this effect applies to both GM and WM, it may disproportionately obscure negative WM responses by bringing them toward zero.

### 4.4 Potential Mechanisms for Negative CBF Responses in the WM

A prominent theory consistent with our findings of negative WM CBF responses is the vascular “steal” or flow-redistribution effect. This theory proposes that during hypercapnia, vascular territories with stronger vasodilatory capacity (i.e. GM) undergo larger and more rapid reductions in vascular resistance, thereby diverting blood flow away from regions with weaker or slower vascular responses (i.e. WM) (Duffin et al., 2018; Sobczyk et al., 2014).

Temporal delays in WM vascular responses relative to GM have also been reported in earlier works (Bhogal et al., 2015; Hartmann et al., 2021). During early hypercapnic stages, preferential GM vasodilation could transiently increase the resistance differential between GM and WM, promoting redistribution effects. As hypercapnia progresses and WM vessels dilate further, this imbalance may diminish. Such dynamics also support the observed strong positive inter-ROI ΔCBF associations (Figure 5c, 5d), and could potentially explain the inter-individual difference observed in this study, as they could be at different stages of coping with the increased CO_2_.

While the redistribution effect has often been associated with vascular pathology such as subarachnoid stenosis and leukoaraiosis, suggesting negative CVR could arise from weakened vascular tone (Duffin et al., 2018; Mandell et al., 2008; van Grinsven et al., 2024), the presence of negative ΔCBF responses in young healthy individuals suggests that vascular steal could be part of normal vascular physiology. Thus, caution is warranted when interpreting such effects as pathological markers. Nonetheless, the physiological significance and functional consequences of these dynamics remain open questions.

### 4.5 Baseline CBF and Hemodynamic Capacity

We further posited that the inter-individual variability in CBF response could be explained by each individual’s baseline CBF and hemodynamic capacity. That is to say, baseline perfusion and vascularization could also determine the individual’s CBF reactivity. Indeed, a negative association between %ΔCBF and CBF_0_ was observed in most tissue layers (Figure 5a). This finding is in line with the strong negative relationship previously reported between baseline regional CBF and task-evoked %ΔCBF by Kastrup et al. (Kastrup et al., 1999). However, Kastrup et al. also showed that baseline CBF was not associated with quantitative ΔCBF, suggesting that the observed negative association for %ΔCBF may be partially influenced by mathematical coupling, as variability in the denominator (CBF_0_) can intrinsically amplify negative relationships. In contrast, weaker but non-zero slopes were observed for quantitative ΔCBF in the present study (Figure 5b), this likely suggests that mathematical effects alone are insufficient to explain the findings. Physiological mechanisms may therefore additionally contribute to the observed inter-individual variability in the perfusion response. For example, individuals with higher baseline perfusion may exhibit partial vascular pre-dilation or limited vascular reserve capacity (Halani et al. 2015), thereby limiting further vasodilatory responses during hypercapnia. At the same time, greater baseline perfusion may also provide a larger baseline blood pool available for redistribution, potentially facilitating the vascular steal effects in regions with weaker vasodilatory capacity.

### 4.6 Limitations and Future Directions

Although our WM CBF estimation demonstrated robustness with sufficient data inclusion, potentially prolonged ATT beyond the maximum PLD may still introduce systematic biases in CBF estimation at some very deep WM regions. Furthermore, the hypercapnia-induced ATT shortening may result in residual gas condition-dependent bias in CBF quantification. Further investigation is necessary to systematically evaluate the influence of various within-voxel arrival times on a regional basis under different physiological conditions.

The tissue-dependent heterogeneity of hypercapnic responses observed here has implications for multimodal MRI investigations. Hypercapnia is frequently employed as a vascular modulator in diffusion MRI studies under the assumption of spatially uniform hemodynamic effects (Ding et al., 2012; Miller et al., 2007). The present findings highlight the necessity of further investigating the perfusion-diffusion interactions, with the spatial and temporal complexity of hemodynamics incorporated.

## 5 Acknowledgements

The authors would like to acknowledge financial support from the Natural Sciences and Engineering Research Council of Canada (NSERC), the Canada Research Chairs Program (JJC), and funding support from Ydessa Hendeles Graduate Scholarship (YLS). We extend our special thanks to Dr. Colette Milbourn for designing and developing the gas delivery system used in this study. We are also grateful to all participants for their time and commitment to this research.

## 6 Data and Code Availability

The code will be made available upon request. The data cannot be shared publicly due to ethics approval restrictions and can be shared with explicit approval from the research ethics board. Data and code can be accessed by contacting the author YLS or JJC.

## 7 Author Contributions

**Yutong Lydia Sun:** Conceptualization, data curation, formal analysis, investigation, methodology, visualization, writing - original draft preparation. **Nayana Menon:** Data curation, formal analysis, investigation, writing - review and editing. **Xiaole Zachary Zhong:** Software, formal analysis, writing - review and editing. **J. Jean Chen:** Conceptualization, funding acquisition, investigation, methodology, project administration, supervision, writing—review and editing.

## 8 Declaration of Competing Interests

The authors declared no potential conflicts of interest with respect to the research, authorship, and/or publication of this article.

## 9 Ethics

The study was approved by Baycrest REB (#11-47).

## 11 Supplementary Material

**Figure 1.**
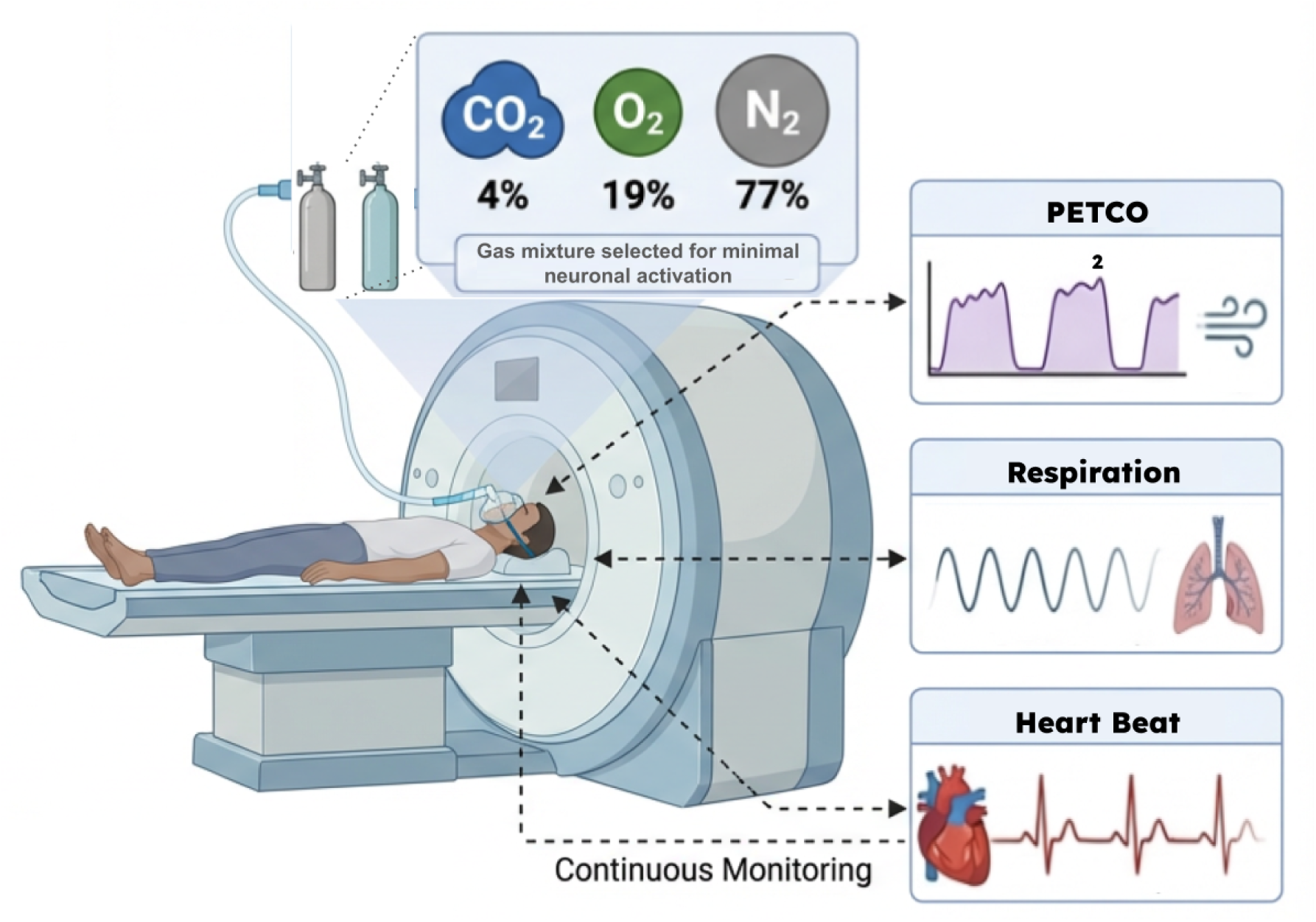
Experimental setup for controlled hypercapnia during MRI acquisition with physiological monitoring. Participants underwent MRI scanning while breathing a controlled gas mixture consisting of 4% CO₂, 19% O₂, and 77% N₂, delivered via a custom gas administration system. The gas delivery apparatus was connected to the participant through a mask to ensure stable and reproducible inhalation throughout the scan. Continuous physiological monitoring was performed, including end-tidal partial pressure of CO_2_ (PETCO₂), respiratory rate, and heart pulse, to verify effective induction of hypercapnia and to ensure participant safety and physiological stability during the experiment.

**Figure 2.**
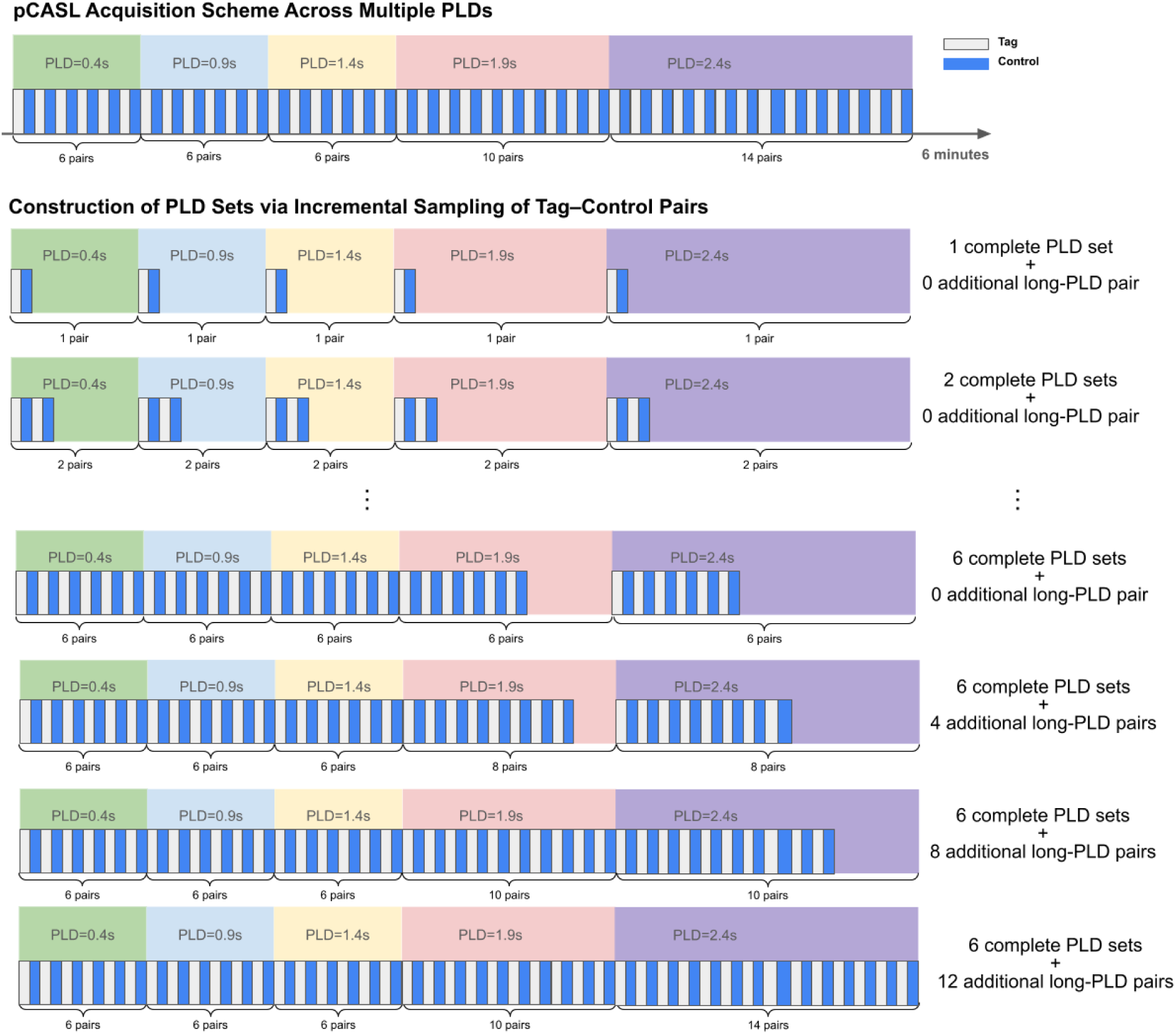
Multi-PLD pCASL acquisition scheme and construction of PLD sets for analysis. Top: Schematic of the full pCASL acquisition spanning approximately 6 minutes, illustrating the sequential acquisition of tag–control pairs across five post-labeling delays (PLDs: 0.4, 0.9, 1.4, 1.9, and 2.4 s). Each PLD block contains a predefined number of tag–control pairs, with longer PLDs sampled more densely to improve signal stability in regions with prolonged arterial transit times. Bottom: Illustration of the procedure used to construct PLD sets by incrementally extracting tag–control pairs from each PLD. Starting from a minimal dataset containing one pair per PLD (i.e., one complete PLD set), additional pairs are progressively included to generate increasingly larger datasets. This approach enables systematic evaluation of data quantity for CBF and ΔCBF estimations.

## Notes

### Competing Interest Statement

The authors have declared no competing interest.

### Summary of Updates

This version has been substantially revised following preparation for journal submission. The manuscript has been reorganized to improve clarity, with a more focused Introduction that better distinguishes the motivations for CBF-based versus BOLD-based cerebrovascular reactivity studies and more clearly defines the study objectives. The Methods section has been expanded to provide additional methodological detail, including software versions, MRI acquisition parameters, ROI generation procedures, morphological operations used to define tissue-depth-specific regions, quality-control procedures, and statistical analyses. The Discussion has been extensively revised to strengthen the physiological interpretation of the findings, including a more comprehensive discussion of white-matter perfusion quantification, arterial transit time, technical limitations of ASL, the distinction between BOLD- and CBF-based cerebrovascular responses, and the potential roles of vascular resistance, vascular reserve capacity, and flow redistribution in explaining heterogeneous white-matter responses. The interpretation of the main findings has also been refined to emphasize that the results challenge the assumption of uniformly positive cerebral blood flow responses during hypercapnia, while distinguishing this from an overall increase in global cerebral blood flow. Additional references have been incorporated throughout the manuscript to better contextualize the findings within the existing literature. Figure legends, terminology, and text have been revised for improved consistency and readability, and numerous minor edits were made to grammar, formatting, and scientific wording. These revisions improve the clarity and completeness of the manuscript without altering the underlying data, analyses, or principal conclusions.

